# Conditional *Dystrophin* ablation in the skeletal muscle and brain causes profound effects on muscle function, neurobehavior, and extracellular matrix pathways

**DOI:** 10.1101/2025.01.30.635777

**Authors:** Muthukumar Karuppasamy, Katherine G. English, James R. Conner, Shelby N. Rorrer, Michael A. Lopez, David K. Crossman, Jodi R. Paul, Miguel A. Monreal-Gutierrez, Karen L. Gamble, Karyn A. Esser, Jeffrey J. Widrick, Louis M. Kunkel, Matthew S. Alexander

## Abstract

Duchenne muscular dystrophy (DMD) patients suffer from skeletal and cardiopulmonary weakness, and interestingly up to one third are diagnosed on the autism spectrum. Dystrophin is an essential protein for regulating the transmission of intracellular force to the extracellular matrix within the skeletal muscle, but also plays key roles in neurobehavior and cognitive function. The mouse dystrophin gene (also abbreviated *Dmd*) is X-linked and has several isoforms with tissue-specific expression, including the large *Dp427m* muscle transcript found in heart and skeletal muscle, and the *Dp427c* transcript that encodes the brain-specific dystrophin cerebellar protein. Understanding the functional requirements and pathways that are affected by dystrophin loss will impact dystrophin replacement gene therapy and exon-skipping correction strategies. We generated conditional *Dystrophin* knockout mice by targeting exon 52 of the mouse *Dystrophin* (*Dmd*^flox52^) locus. We generated dystrophin constitutive and inducible myofiber knockout (*Dmd* mKO) mice to evaluate the tissue-specific function of the large skeletal muscle dystrophin isoform. Constitutive embryonic deletion of the *Dystrophin* gene exclusively in skeletal myofibers resulted in a severe skeletal muscle myopathy, dystrophic histopathology, and functional deficits compared to the *mdx* mouse. Transcriptomic analysis of skeletal myofibers of the *Dmd* mKO mice revealed the dysregulation of key extracellular matrix and cytokine signaling pathways. Separately, we generated Purkinje neuron cerebellar dystrophin knockout (*Dmd*:Pcp2 KO) mice that displayed neurobehavioral deficits in social approach, social memory, and spatial navigation and working memory. These studies reveal the essential requirement for dystrophin expression in both the skeletal muscle and brain for normal physiological and neurobehavioral function.

**Significance Statement:** Duchenne muscular dystrophy is caused by the lack of a functional dystrophin protein in muscle. The large dystrophin (Dp427m) isoform is expressed in skeletal, cardiac, and smooth muscle, but its tissue-specific requirements remain unknown. We generated and characterized a conditional skeletal muscle knockout mouse (*Dmd* mKO). Constitutive embryonic genetic ablation of skeletal muscle *Dystrophin* resulted in muscle histopathologies similar to the *mdx* mouse, while postnatal muscle *Dystrophin* ablation resulted in milder pathologies. Ablating cerebellar *Dystrophin* Dp427c using a *Pcp2/L7*-Cre driver resulted in sociobehavioral defects. Transcriptomic analysis of the *Dmd* mKO mice showed a severe reduction of extracellular matrix and cytokine signaling pathways. Our study reveals an essential role for skeletal muscle dystrophin and identifies essential pathways for modulation using dystrophin-replacement therapies.

## Introduction

In Duchenne muscular dystrophy (DMD) patients there is a disruption of the dystrophin gene (also abbreviated *DMD*) that is predominantly expressed in skeletal and cardiac muscle. DMD patients develop progressive muscle weakness in their teenage years that results in severe muscle degeneration and cardiorespiratory complications. The *DMD* gene is X-linked, thus DMD predominantly affects males who lose the ability to walk by their early teenage years. Mammalian dystrophin protein is an essential protein that is required for normal skeletal muscle function through its key binding to the actin cytoskeleton and dystrophin-associated protein complex (DAPC). When fully sequenced, the human *DMD* locus was found to be extremely large as it spans 2.2 megabases (Mbs) and has multiple coding regions that are essential for the function of the dystrophin protein(1). The regulation of the dystrophin protein is complex in that the major protein (Dp427) is expressed in multiple tissues and organs including skeletal muscle, heart, lung, intestine, and the brain(2). Further complexity of dystrophin transcriptional regulation exists in that multiple isoforms and *DMD* promotors exist and the regulation of its expression on a tissue-specific manner has yet to be fully elucidated.

The most widely used Duchenne mouse model is the *mdx* mouse, which harbors an *Dmd* exon 23 nonsense mutation (C>T) at nucleotide 3185 of the large *Dp427m* RNA transcript(3, 4). Several other N-ethyl-N-nitrosourea (ENU) mutagenesis and targeted knockout mouse strains have been generated since the identification of the *mdx* mouse, that all show varying muscle phenotypes based on the DMD mutation type and *Dystrophin* exon(s) targeted(5). Additional DMD mouse and other animal models have been generated and assessed for their dystrophin expression, mutation location, muscle pathophysiologies, and their ability to respond to pre-clinical and clinical corrective drugs. Attempts to generate conditional mouse models have had mixed outcomes due the large *DMD* gene loci and difficulties in targeting the X chromosome using gene editing. One study demonstrated that a dual *Dmd* exon 2 and 79 floxed mouse when mated with a tamoxifen-inducible cardiac and skeletal muscle *MerCreMer* transgene resulted in 25-35% dystrophin gene excision(6). A previously homologous recombination (HR)-established knockout mouse targeting dystrophin exon 52 (a known DMD mutation “hot spot” region) resulted in the disruption of the large Dp427, Dp260, and Dp140 dystrophin isoforms(7). This dystrophin exon 52 model (*mdx52*) displayed severe muscle degeneration pathologies, in addition to cardiac and neurobehavioral defects similar to those observed in the *mdx* (*Dmd* exon 23) mouse(7–10). More in-depth comparisons of multiple DMD mouse strains have revealed neurological, cognitive, and behavioral deficits compared to WT controls(11, 12). Using this knowledge, we generated a conditional-ready *Dmd* mouse by inserting loxP sites flanking *Dystrophin* exon 52 (*Dmd^flox52^*) that allowed for the ablation of the large *Dp427* isoforms upon Cre-mediated excision. Combinatorial mating with skeletal myofiber Cre lines allowed us to define the tissue-specific requirements of the dystrophin protein in skeletal muscle in early and post-natal muscle growth and regeneration. Separate matings with a Purkinje neuronal Cre line also recapitulated several of the learning, cognitive, and social deficits as other DMD mouse lines.

## Results

### Generation and initial characterization of skeletal muscle Dystrophin knockout mice

We performed CRISPR-targeting of the mouse *Dystrophin* exon 52 locus and inserted two loxP sequences in the intronic region immediately flanking the exon (**Figure 1A**). Upon Cre-mediated excision of *Dmd* exon 52, the resulting DNA sequence contains an out-of-frame deletion that impairs the stability of the Dp427, Dp260, and Dp140 transcripts as has been previously shown in the *mdx52* mouse(13). We mated male *Dmd^flox52^*^/Y^ mice with the constitutive human skeletal actin (*HSA*)-Cre to ablate expression of the large Dp427m protein within the skeletal myofibers (**Figure 1B**). We compared our conditional dystrophin knockout mouse model to the *mdx^5cv^* mouse that has well-characterized dystrophic histological, molecular, and physiological muscle symptoms(14). The resulting *Dmd^flox52^*^/Y^:HSA-Cre+ (*Dmd*:HSA KO) mice have no overt pathologies at birth, but similar to many *mdx* strains develop a progressive muscle degeneration starting between 4-6 months of age (**Figure 1C**). Analysis of both fast twitch *Tibialis anterior* (TA) and slow twitch (soleus) muscles from *Dmd*:HSA KO mice showed increased centralized myonuclei, large areas of fibrosis, and increased collagen production throughout the myofibers (**Figure 1C**). *Dmd*:HSA KO diaphragm muscle also revealed hallmark muscle degeneration, increased centralized myonuclei, and the presence of collagen deposits compared to Cre-controls (**Figure 1C**). Cross-sectional area (CSA) of *Dmd*:HSA KO TA myofibers showed an increased frequency of smaller myofibers compared to Cre-controls and similar to those found in our *mdx^5cv^* mice (**Figures 1D and 1E**). Quantitative western blot assessments of Dp427m protein levels in the *Dmd*:HSA KO TA and soleus muscles showed an average of <5% remaining Dp427m protein in the TA and the soleus muscles (**Figure 1F**). This was further confirmed using immunofluorescent antibody staining in these muscles showing severely reduced levels of the large Dp427m protein (**Supplemental Figure S1**).

**Figure 1.**
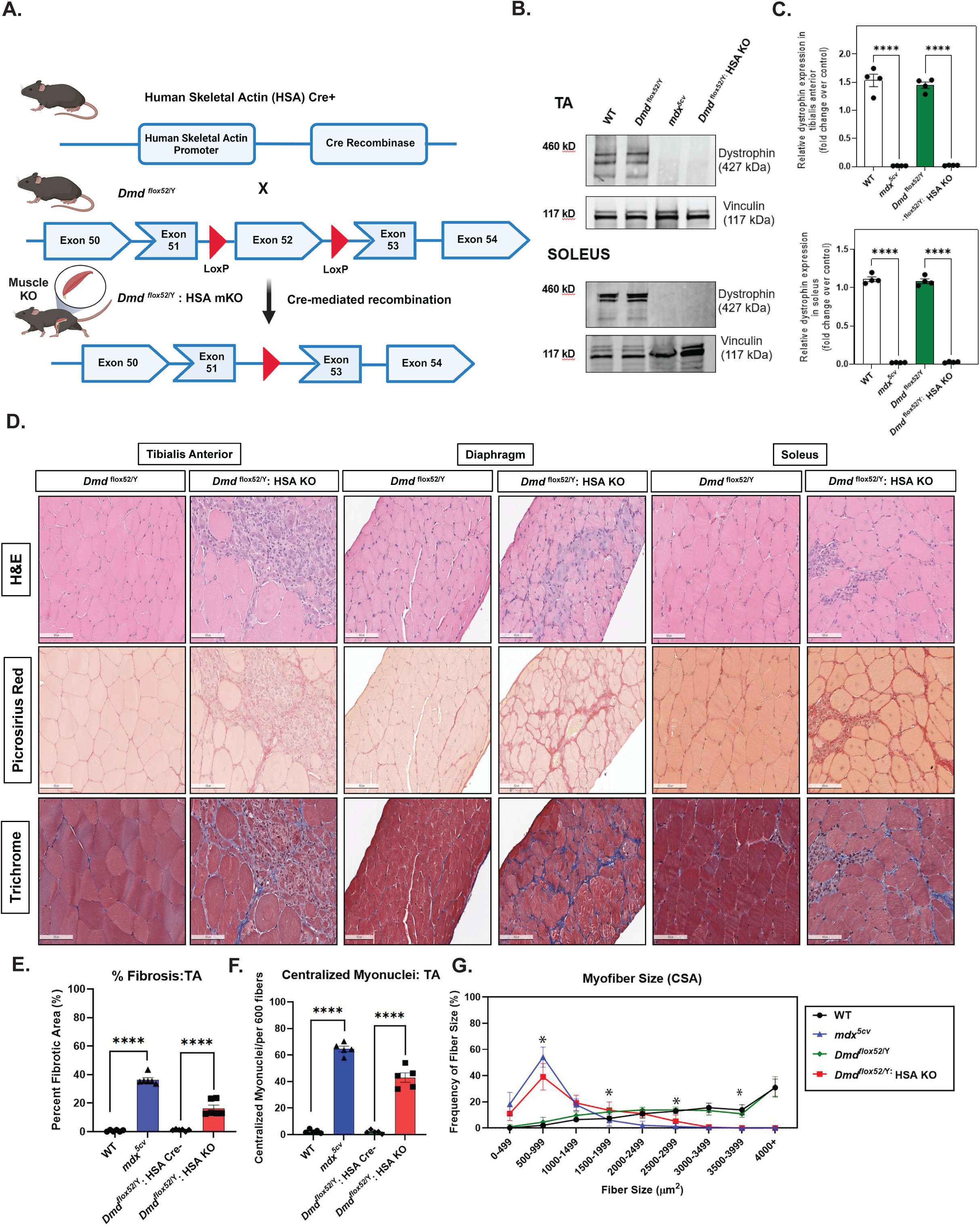
Characterization of a myofiber-specific *Dmd* conditional knockout mouse. **A.** Schematic outlining the generation of myofiber-specific Dystrophin conditional knockout mice (*Dmd*:HSA KO) by breeding our *Dmd^flox52^*^/Y^ mice to human skeletal actin (*HSA*)-Cre transgenic mice. Cre-mediated excision of *Dmd* exon 52 results in an out-of-frame disruption and instability of the Dp427 RNA transcript. **B.** Western immunoblotting for the large Dp427m protein in the TA and soleus muscles of WT (open), *Dmd^flox52^*^/Y^ (blue), *mdx^5cv^* (red), and *Dmd*:HSA KO (green) adult (6-month-old) male mice. **C.** Densitometry graphs WT, *Dmd^flox52^*^/Y^, *mdx^5cv^*, and *Dmd*:HSA KO immunoblots taken from TA and soleus muscle lysates and normalized to Vinculin protein. N = 5 separate mice per cohort. **D.** Histochemical staining of TA, soleus, and diaphragm muscles isolated from the WT, *Dmd^flox52^*^/Y^, *mdx^5cv^*, and *Dmd*:HSA KO male mice. Hematoxylin and eosin (H&E), picrosirius red, and Masson’s trichrome histochemical staining performed. Scale bar = 100 µm. **E.** Percent fibrosis (%) in the TA muscles of the four mouse cohorts analyzed. **F.** Percentage (%) centralized myonuclei for the approximately 600 myofibers of the TA muscles of the four mouse cohorts analyzed. **G.** Cross-sectional area (CSA) displaying myofiber sizes (µm^2^) shown for each of the four cohorts. Two-way ANOVA with Bonferroni correction performed. P-values of significance are shown: * p < 0.05, ** p < 0.01, *** p < 0.005, and **** p < 0.001.

### Skeletal myofiber loss of muscle dystrophin greatly impairs muscle physiological function

We performed a muscle functional analysis of the *Dmd*:HSA KO adult mice by performing an assessment of muscle performance following TREAT-NMD standard operating procedures (SOPs) for *mdx* mice(15). *Dmd*:HSA KO mice showed reduced rotarod latency to fall, open field test (OFT) total distance travelled, and impaired forelimb grip strength (**Figures 2A-2C**). All of these locomotor and muscle strength metrics were comparable to those of aged-matched *mdx* mice. *Dystrophin*-deficient mice have impaired treadmill running performance due in part to the lack of sarcolemma-associated neuronal NOS (nNOS) binding to Dp427 at the spectrin-like repeats encoded by *Dmd* exons 45-48(16, 17). We performed a standardized treadmill downhill running assessment of age-matched male WT, *mdx^5cv^*, *Dmd^flox52^*^/Y^ (Cre negative), and *Dmd*:HSA KO mice. The *Dmd*:HSA KO and *mdx^5cv^* mice performed similarly in showing quicker times to exhaustion, reduced overall distance travelled, and reduced velocity compared to internal cohort control mice (**Figures 2D-2F**). These findings are consistent with the requirement of nNOS-binding to skeletal muscle dystrophin as essential for skeletal muscle exercise and running performance(18, 19). We assessed physiological force in EDL muscles from our *Dmd*:HSA KO mice to determine if there were functional deficits in the peak force and resistance to high mechanical strain. Despite their greater mass (**Supplemental Figure S2A**), EDL muscles from the *Dmd*:HSA KO mice attained similar peak force as EDL’s from the *Dmd^flox52^*^/Y^ Cre-controls (**Supplemental Figure S2B**). This reduction in force per unit physiological cross-sectional area of muscle (**Supplemental Figure S2C**) indicates a reduction in the quality of the *Dmd*:HSA KO EDL. Eccentric contractions reduced force of muscles from both groups of mice but this response was exacerbated in the *Dmd*:HSA KO mice (**Supplemental Figures S2D and S2E**). These results suggest that skeletal muscle Dystrophin loss has a moderate reduction in myofiber force output and resistance to mechanical strain in comparison with that of *mdx* mice.

**Figure 2.**
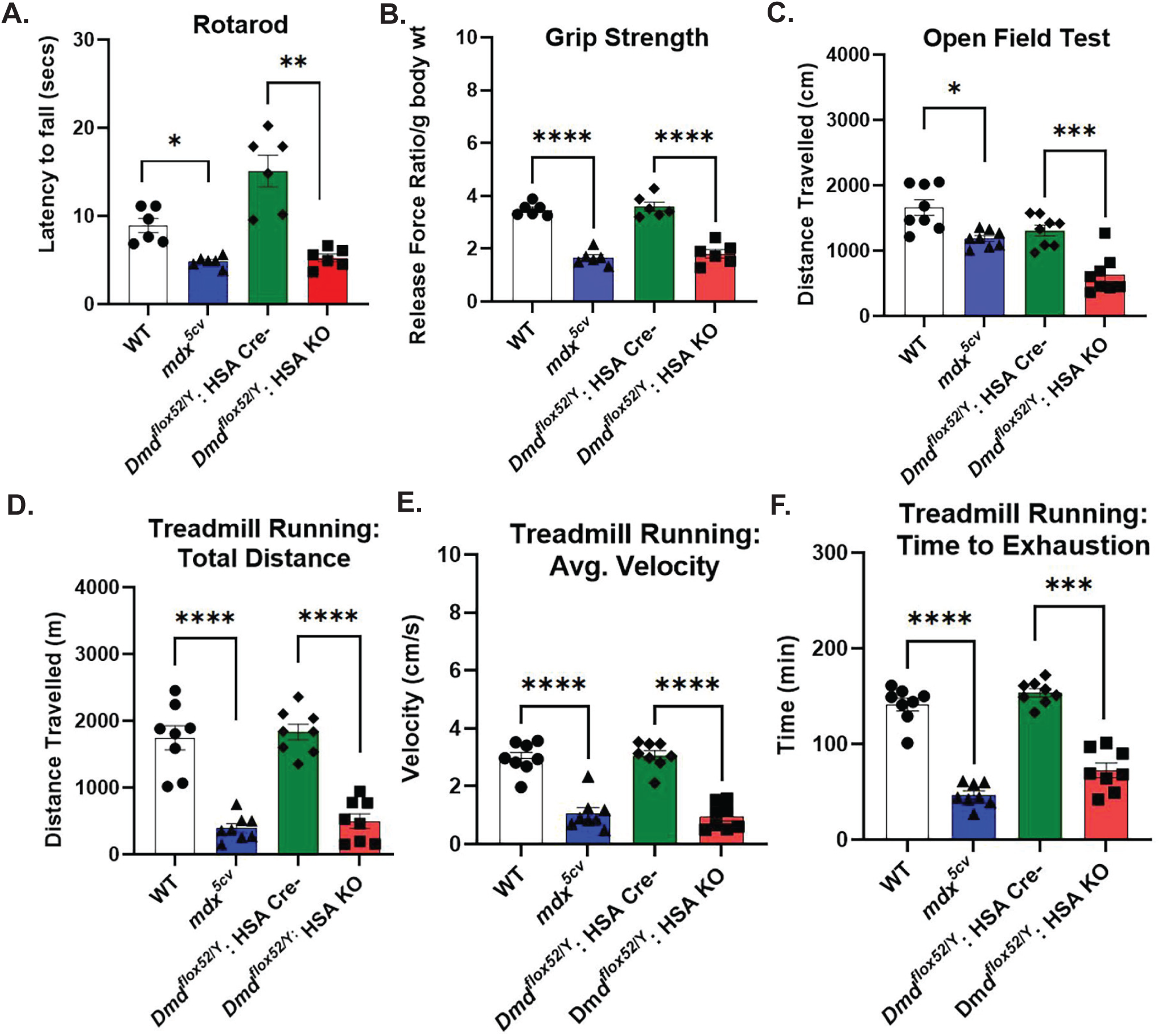
Adult *Dmd* myofiber KO mice have a poor running capacity. **A.** Rotarod analysis shown for the four mouse cohorts: WT, *Dmd^flox52^*^/Y^, *mdx^5cv^*, and *Dmd*:HSA KO. **B.** Grip strength measured for mouse forelimbs shown normalized to grams (g) body weight. **C.** Open field testing (OFT) of the four cohorts of mice showing distance traveled in centimeters (cm). **D.** and **E.** Forced downhill treadmill running of the four cohorts of mice with average velocity and time to exhaustion shown. N = 6 mice/cohort. Two-way ANOVA with Bonferroni correction performed. P-values of significance are shown: * p < 0.05, ** p < 0.01, *** p < 0.005, and **** p < 0.001.

### *Dystrophin* myofiber KO mice have impaired skeletal muscle regeneration following injury

We performed a cardiotoxin (ctx)-induced injury to the TA muscles of the *Dmd*:HSA KO and control *Dmd^flox52^*^/Y^ mice to evaluate the regenerative capacity of the skeletal muscle in the absence of the Dp427m protein. The *Dmd*:HSA KO mice showed a significant amount of fibrosis, collagen, and necrotic regions within the muscle that was comparable to the *mdx^5cv^* strain (**Supplemental Figure S3A**). Immunofluorescent analysis of the myofibers from the *Dmd*:HSA KO mice showed elevated levels of centralized myonuclei after 21 days post ctx-injury (**Supplemental Figure S3B**). These results demonstrate the absolute requirement for the dystrophin protein in the myofiber and suggest that dystrophin from the muscle satellite cells (MuSCs) cannot compensate for the lack of myofiber dystrophin in the normal regenerative process.

### Skeletal muscle-specific *Dystrophin* loss results in cytokine and ECM pathway disruption

We performed RNA-sequencing on the *Dmd*:HSA KO and Cre-control TA muscles to determine what transcript and signaling pathways were disrupted upon the loss of skeletal muscle *Dystrophin* (**Figure 3A**). Ingenuity Pathway Analysis of RNA transcripts revealed several inflammatory cytokine transcripts that were disrupted as well as extracellular matrix (ECM) including several *Collagen* isoforms (**Supplemental Figure S4**). Interestingly, several transcripts known to be disrupted in DMD patients were also disrupted in the *Dmd* HSA KO mice including an elevation of inducible nitric oxide synthase 2 *(iNos*; *Nos2*) (**Figure 3D**). We noted a strong downregulation in key immune-responsive cytokines including *Tgfb1* which drives DMD muscle pathologies in *Dmd* HSA KO mouse muscles and indicates disruption of key cell-cell signaling between the skeletal myofibers and the immune cells residing in the muscle (**Figures 3B-3D**). These transcriptomic analysis confirms and delineates an essential transcriptomic regulation of key signaling pathways that are known regulators of dystrophin pathophysiology.

**Figure 3.**
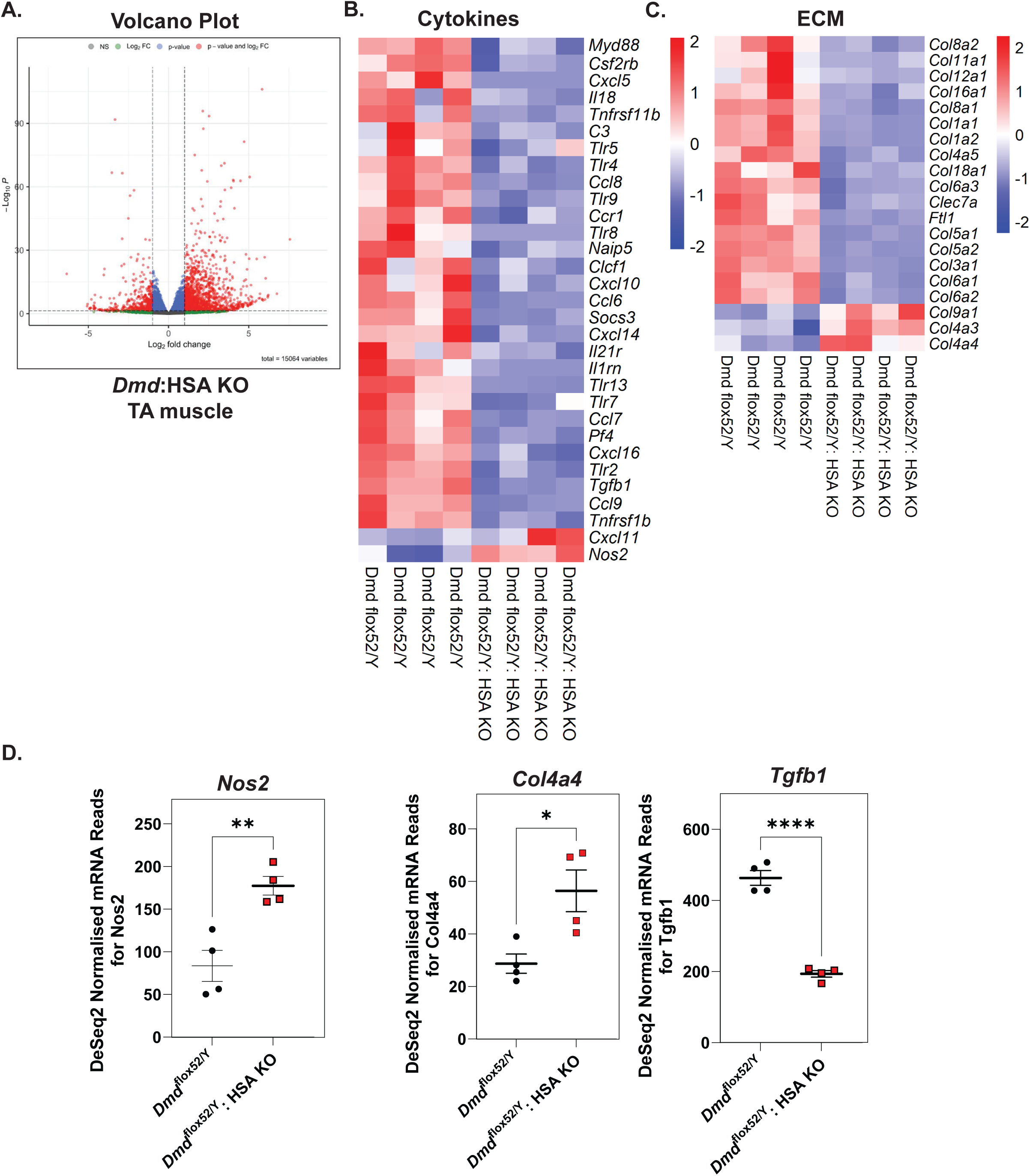
*Dmd* myofiber KO muscles have disrupted cytokine and ECM signaling pathways. **A.** Volcano plot of the approximately 6,200 dysregulated DEGs in the TA muscles of *Dmd*:HSA KO mice compared to *Dmd^flox52^*^/Y^ (HSA-Cre-negative) aged-matched littermate controls. **B.** and **C.** Heatmaps of significantly dysregulated showing significantly elevated (> 2.0 fold) or downregulated (< 2.0 blue) DEGs with an emphasis on cytokine and ECM pathways. D. RNA-seq DeSeq2 FKPM counts of three candidate highly disrupted transcripts that are validated DMD muscle biomarkers: *Nos2*, *Col4a4*, and *Tgfb1* are shown. Four mice of each cohort (*Dmd^flox52^*^/Y^-black; *Dmd*:HSA KO-red) were analyzed for differentially expressed genes (DEGs). Student’s t-test, two tailed used for comparisons. P-values of significance are shown: * p < 0.05, ** p < 0.01, *** p < 0.005, and **** p < 0.001.

### Postnatal skeletal myofiber *Dystrophin* loss impairs muscle maintenance despite dystrophin protein stability

We next sought to evaluate the role of skeletal muscle dystrophin in the regulation of adult skeletal muscle maintenance and regeneration. Subsequently, we mated our conditional *Dmd^flox52^*^/Y^ mice to the *HSA*-MerCreMer tamoxifen-inducible skeletal myofiber Cre line to temporally ablate Dp427m expression in the adult mice (**Figure 4A**). Two months after tamoxifen dosing, we observed 25-30% reduction of total Dp427m protein within the TA skeletal muscles of *Dmd*:HSA-MerCreMer+ (+ Tam) of *Dmd*:HSA-MerCreMer+ (-Tam) controls (**Figure 4B**). We observed a similar 20-30% decrease of total Dp427m protein within the gastrocnemius and soleus muscles in the *Dmd*:HSA-MerCreMer+ (+ Tam) mice (**Figure 4B**). These results are similar to the other reported conditional *Dmd* mice that showed partial reduction of Dp427mprotein and highlights the stability of the large dystrophin protein after 2 months of protein turnover(6). A histological examination of the *Dmd*:HSA-MerCreMer+ (+ Tam) mice showed minimal myofiber degradation and small pockets of regenerating myofibers compared to controls (**Figure 4C**). Overall myofiber cross-sectional area (CSA) and minimal Feret diameter analysis revealed smaller clusters of myofibers in the *Dmd*:HSA-MerCreMer+ (+ Tam) mice compared to no tamoxifen mock controls (**Figures 4D and 4E**). These findings implicate the large amount of stability of the large dystrophin Dp427m protein and implicate protein turnover as an essential part of dystrophin-restoration therapies.

**Figure 4.**
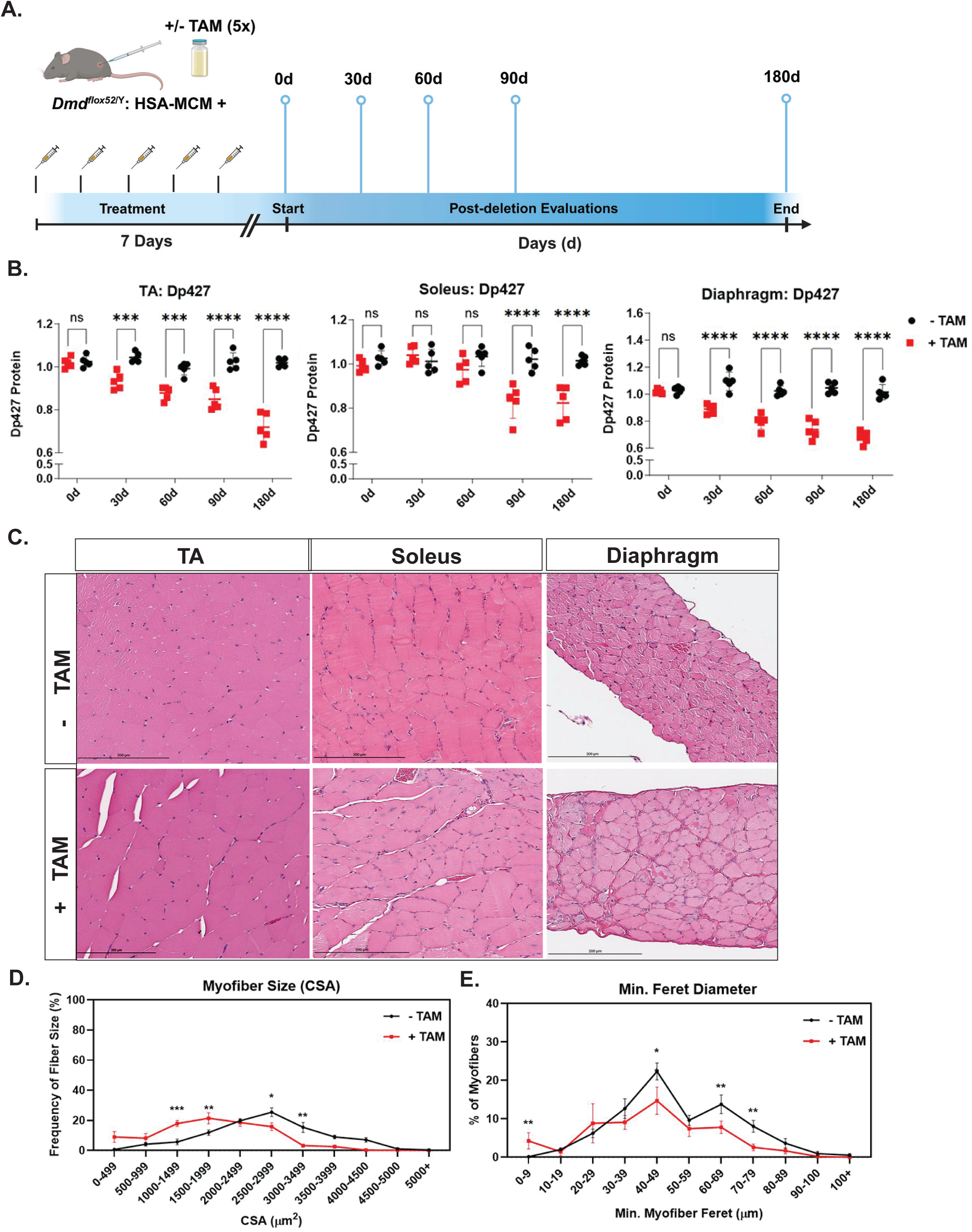
Postnatal loss of muscle dystrophin results in mild skeletal muscle pathologies. **A.** Schematic for conditional ablation of postnatal dystrophin using a tamoxifen-inducible (± TAM) strategy. *Dmd^flox52^*^/Y^ mice were mated to the *HSA*-MerCreMer (HSA-MCM) to conditionally ablate dystrophin (+ TAM; red). *Dmd*:HSA (Cre+) mice given sunflower seed oil (-TAM) were used as controls (black) After a one-week (5x per week) tamoxifen dosing (80 mg/kg body weight) in 8-week-old male mice via intraperitoneal injection, mouse cohorts were evaluated at 0 (pre-dosing), 30, 60, 90, and 180 days (d) post deletion. N = 5 mice per cohort and time period for biopsy. **B.** Densitometry western immunoblot measurements of the large dystrophin (Dp427m) protein isoform measured in the TA, soleus, and diaphragm muscles. Note the robust persistence of the large dystrophin protein and low protein turnover in the skeletal muscle tissues. **C.** H&E evaluation of *Dmd^flox52^*^/Y^:HSA-MCM (Cre+) mice (± TAM) evaluated at 180 days post ablation. Scale bar = 200 µm. **D.** and **E.** CSA and minimum Feret diameter measured at 180 days in the TA muscles reveal smaller groupings of myofibers in the *md^flox52^*^/Y^:HSA-MCM (Cre+) mice (+ TAM) mice compared to (-TAM) controls. Student’s t-test, two tailed used for comparisons. P-values of significance are shown: * p < 0.05, ** p < 0.01, *** p < 0.005, and **** p < 0.001.

### Loss of cerebellar dystrophin impairs neurobehavioral function and results in ASD-like behaviors

Previous studies from our lab and others in various *mdx* strains demonstrated that the Dp427 cerebellar dystrophin isoform was essential for normal neurobehavior and social development(12, 20–22). We mated the conditional *Dmd^flox52^*^/Y^ mice to *L7/Pcp2*-Cre (*Dmd*:Pcp2 KO) mice to ablate the Dp427c protein in Purkinje neurons of the cerebellum (**Figure 5A**). Isolated cerebellums from adult *Dmd*:Pcp2 KO mice showed loss of approximately 65% of cerebellar dystrophin protein compared to Cre- controls (**Figure 5B**; **Supplemental Figure S5**). We performed a series of neurobehavioral assessments in the *Dmd*:Pcp2 KO mice including social approach and social novelty between novel and familiar objects and mice (**Figure 5C**). *Dmd*:Pcp2 KO showed impaired social novelty in object recognition compared to *Dmd^flox52^*^/Y^ Cre- control mice (**Figures 5D and 5E**). We performed a similar 3-chamber assessment (novel mouse versus familiar mouse) and the *Dmd*:Pcp2 KO mice showed no preference for either mouse as compared to controls suggesting that social behavior defects are linked to cerebellar dystrophin protein expression. We performed both T- and Y-maze assessments in the *Dmd*:Pcp2 KO mice and found no differences in T-maze alternations compared to Cre- control mice (**Figure 5F**). However, in the Y-maze assessment, *Dmd*:Pcp2 KO mice had reduced alternations without a change in total arm visits which together suggest reduced spatial working memory compared to Cre- controls (**Figure 5G**). Memory tests were also conducted on the *Dmd*:Pcp2 KO mice that displayed impaired time to find the platform in Morris water maze (**Figure 5H**). These findings support an essential role for cerebellar Purkinje neuronal expression of *Dystrophin* and demonstrate a functional role for Dp427 in the regulation of neurobehavior in DMD.

**Figure 5.**
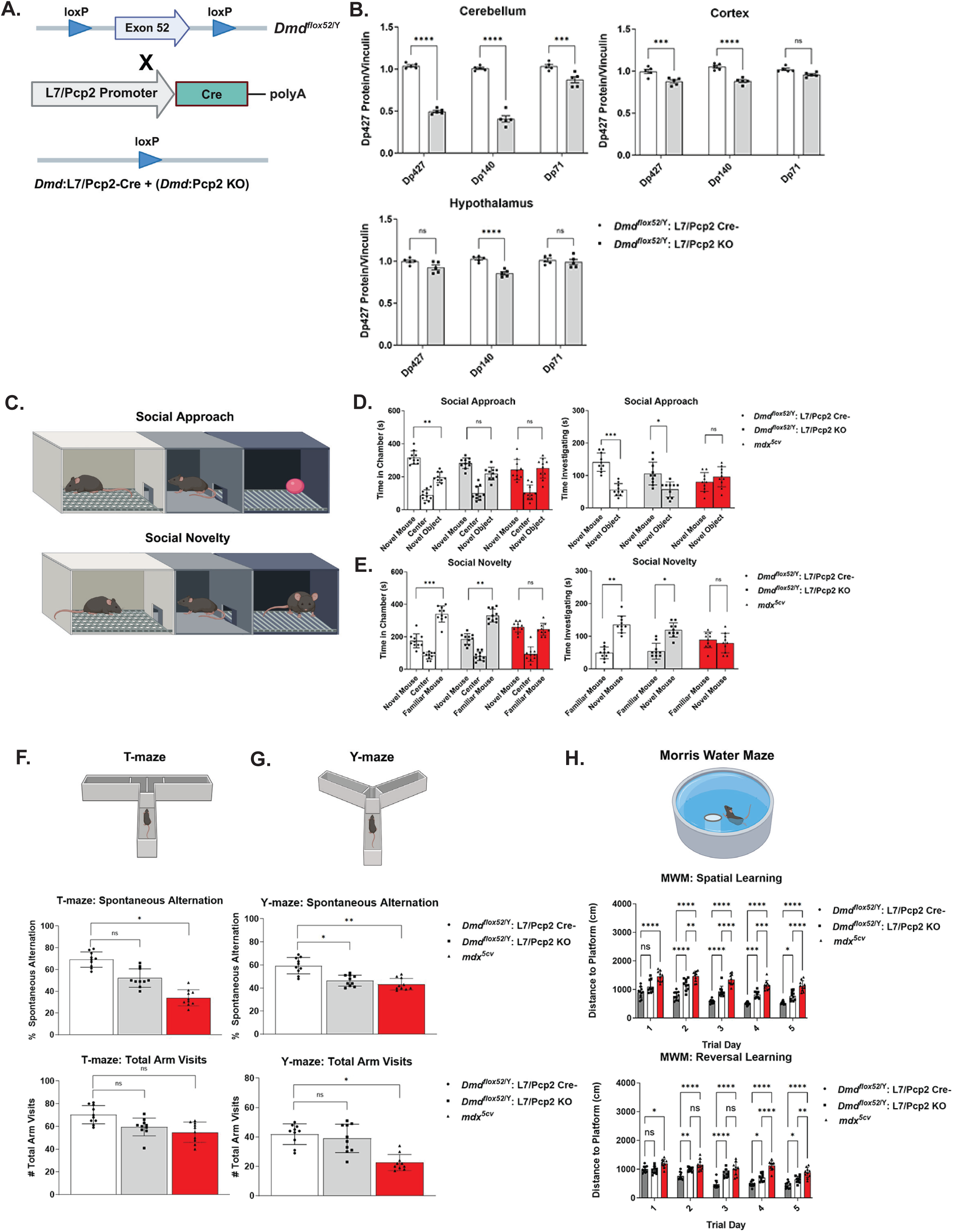
Loss of Purkinje neuron brain *Dystrophin* impairs neurobehaviors in mice. **A.** Breeding schematic for the *Dmd^flox52^*^/Y^ line to the *L7/Pcp2*-Cre (cerebellar Cre driver) transgenic line to generate Dmd:Pcp2 KO mice. **B.** Densitometry western immunoblot quantification of the large dystrophin (Dp427) protein expression in the cerebellum, cortex, and hypothalamus regions of the brain in the *Dmd^flox52^*^/Y^ (Cre-; open) and *Dmd*:Pcp2 KO (Cre+; grey) cohorts. N = 5 mice per cohort were analyzed. **C.** Graphics showing the social approach (novel mouse versus novel object) and social novelty (novel mouse versus familiar mouse) assays performed on the cohorts of mice in a 3-chamber assay system. **D.** and **E.** Social approach and social novelty time in chambers (1, 2, and 3) shown between the 3 analyzed mouse cohorts: the *Dmd^flox52^*^/Y^ (open bars), *Dmd*:Pcp2 KO (grey bars), and *mdx^5cv^* (red bars). Time in chamber and time investigating in minutes (mins) are shown. N = 10 mice per cohort were analyzed with an acclimation period of equivalent amounts. **F.** and **G.** T- and Y-maze assessments of the 3 mouse cohorts are shown with spontaneous alternation and the number of total arm visits. **H.** Morris water maze (MWM) assessments in the 3 cohorts are shown as distance to platform (cm) compared to trial day. Spatial and reverse learning assessments were performed on separate days. N = 10 adult (6-month-old) mice were analyzed for each cohort. Two-way ANOVA with Bonferroni correction performed. P-values of significance are shown: * p < 0.05, ** p < 0.01, *** p < 0.005, and **** p < 0.001.

## Discussion

The generation of novel DMD animal models using CRISPR gene editing and other next-generation genomic DNA editing tools highlights the growing need for comprehensive animal models that accurately reflect disease symptoms and pathophysiologies(23). Our conditional dystrophin knockout model allows for the tissue-specific study of the impact of both dystrophin loss and when combined with dystrophin-replacement strategies, correct DMD strategies. Newer dystrophinopathy mouse models have identified the impact of dystrophin loss on muscle disease symptom progression. The *bmx* (Becker muscular dystrophy; *Dmd* exons 45-47 in-frame deletion) mouse has been useful in elucidating the slower muscle disease progression that occurs in patients(24). Our conditional approach allows for the exploration of dystrophin requirements at key developmental and DMD disease-relevant time points.

Previous attempts to delineate skeletal muscle dystrophin protein expression using AAV-shRNAi strategies have revealed dystrophin as essential for sarcomeric organization within the muscle myofiber(25). A study of cultured *mdx* mouse myoblasts revealed a similar downregulation of Collagen and histone transcripts along with impaired fusion and proliferation markers(26). Indeed, there may be a significant amount of epigenetic remodeling that occurs which was shown in embryonic stem (ES) cell cultures of *mdx* mice and human induced pluripotent stem cells (hiPSCs)(27, 28). Given the alteration of myogenic determinant transcriptional pathways, consideration of the impact of *Dystrophin* loss on epigenetic influences on myogenic differentiation is warranted(29). Key autophagy and PTEN/AKT pathways have been shown to be targets for therapeutic amelioration of dystrophic muscle pathologies(30–32). Questions remain on the impact and influence of the large dystrophin muscle isoform on muscle satellite cells (MuSCs) and its regulation of cellular polarity(33, 34). The Dp427 protein is expressed in muscle satellite cells and it is currently unclear if restoration of dystrophin expression in the MuSC population could compensate for myofiber dystrophin loss. Therapies to restore MuSC polarity in DMD MuSCs may be beneficial for restoring the cellular kinetics and polarity defects observed in DMD muscles(35, 36).

It is evident that the expression of dystrophin protein does occur in the brain and both large and smaller dystrophin isoforms likely play independent and critical neuronal functions. Clinical studies have demonstrated that loss of dystrophin is important for normal neurological and neurobehavioral function as up to one-third of all DMD patients are diagnosed on the ASD spectrum with symptoms ranging from mild cognitive impairment to severe developmental delay(11, 37–39). Evaluation of DMD mouse models have shown impaired neurobehavioral responses in memory, learning, and overall electrophysiological neuronal function(22, 40–42). Our *Dmd*:Pcp2 KO mice show similar defects to those in *mdx* mice, and highlight the essential role for the large brain dystrophin protein expression. These results argue that newer platforms for exon-skipping drug compounds that can more effectively penetrate the blood-brain-barrier (BBB) may be required for the systemic restoration of dystrophin expression(43, 44). Our study demonstrates the absolute requirement for dystrophin expression within the skeletal muscle and brain and warrants further evaluation of the tissue-specific pathways impacted by dystrophin loss. Elucidation of these dystrophin-regulated pathways will aide in dystrophin-replacement and correction therapeutic strategies focused on the amelioration of DMD disease pathologies.

## Materials and Methods

### Mice

Wild type (*C57BL/6J*; strain #000664; Jackson Labs) and *mdx^5cv^* (*C57BL/6J*; strain #002379; Jackson Labs) used in all experiments were maintained in the Alexander lab colony under pathogen-free conditions. Constitutive HSA-Cre (*Acta1*-Cre; strain# 006149) and tamoxifen-inducible *HSA-MerCreMer* (*HSA*-MCM; strain# 031934) were purchased from Jackson Labs (Bar Harbor, ME) and maintained on a congenic *C57BL/6J* background. The *L7/Pcp2*-Cre mice were originally purchased from Jackson Labs and maintained on a congenic *C57BL/6J* background that had been backcrossed greater than 6 generations (strain #006207; Jackson Labs). All mice had a standard Teklad Global Rodent Diet (Inotiv; West Lafayette, IN) with *ad libitum* access to food and water. All mouse strains and protocols were approved by the UAB Institutional Animal Care and Use Committee (IACUC) under the protocol 21393.

### Generation of conditional Dystrophin knockout mice

Conditional *Dystrophin* knockout mice *(Dmd^flox52^*^/Y^) were generated commercially (Cyagen; Santa Clara, CA) via CRISPR guide editing of genomic DNA by inserting loxP sites in the intronic regions flanking mouse *Dystrophin* exon 52 using a 12.2 kb homologous recombination vector. Briefly, loxP sequences 5’-ATAACTTCGTATAGCATACATTATACGAAGTTAT-3’ were inserted in the intronic regions flanking the mouse *Dmd* exon 52 (*Dystrophin* NCBI transcript: NM_007868.6) in *C57Bl/6J* mouse ES cells (mES). Positively-targeted mouse ES cells were screened for neomycin cassette (later removed via FRT-mediated recombination) and diphtheria toxin subunit A (DTA) selection. After PCR based confirmation of the presence of the loxP sites and removal of the neomyocin cassette, the resultant ES cell line was injected into pseudopregnant females. The subsequent F_0_ offspring were genotyped for presence of the loxP sites at mouse *Dystrophin* exon 52, and then positively-targeted males were mated to WT (*C57Bl/6*) females to generate F_1_ offspring. All positively-targeted F_1_ mice were then maintained on the *C57Bl/6* strain for subsequent matings. Mice were genotyped for the *Dmd^flox52^*^/Y^ allele using a forward primer “5-TGACTGCATCTGCATACGTG-3” and reverse “5-GAAGCCTGGTAGGCACCAT-3”. Using the Roche Expand Long Template PCR System (MilliporeSigma; Burlington, MA; Cat# 11681842001) the wild type band (2226 bp), targeted loxP conditional allele (2401 bp), and Cre-mediated excised band (992 bp) were used for genotyping following the manufacturer’s recommendations. Later genotyping was performed commercially using Transnetyx probes (Cordova, TN). Isolation of genomic DNA from TA muscles, cerebellum, and tail DNA were performed using Viagen DirectPCR Lysis Reagent (Viagen Biotech; Los Angeles, CA; Cat# 102-T).

### Histology and Immunofluorescent Imaging

Skeletal muscles were slowly cryofrozen as unfixed tissues in Scigen TissuePlus O.C.T. Compound using an isopentane (FisherScientific; Cat# AC397221000) and liquid nitrogen bath as unfixed tissues. Some muscle biopsies were placed in 30% sucrose (MilliporeSigma; Cat# S0389) sink overnight at 4°C prior to the slow cryofreezing to preserve histological and protein integrity. Muscles were sectioned on a Leica CM1950 cryotome at 7-10 µm thickness, and specimens were placed on Fisherbrand Tissue Path Superfrost Plus Gold slides (Fisher Scientific; Cat# FT4981gplus). H&E staining was performed using Hematoxylin (MilliporeSigma; Cat# H9627) and Eosin Y (MilliporeSigma; Cat# 318906) solutions in a previously described protocol(45). Masson’s trichrome staining was performed using a Masson’s Trichrome Stain Kit (Polysciences, Inc; Warrington, PA; Cat# 25088-1) and picrosirius red staining was performed using a Picrosirius Red Stain Kit (Polysciences, Inc; Cat# 24901-500). For immunofluorescent staining, tissue slides were blocked for one hour in a solution of 10% goat serum/1xPBS and incubated for one hour at room temperature using a M.O.M kit (Vector Labs; Cat# BMK-2202; Newark, CA). Tissue slides were set using a VECTORSHIELD Antifade mounting medium with DAPI kit (Vector Labs; Cat# H-1200-10),

### Immunoblotting

Immunoblotting was performed on protein lysates using tissues that were snap frozen in liquid nitrogen and were homogenized using a bead homogenizer and lysing matrix A beads (MP Biomedicals; Cat# 1169100-CF) following the manufacturer’s protocol. Samples were homogenized in T-PER™ Tissue Protein Extraction Reagent (ThermoFisher Scientific; Cat# 78510) supplemented with a protease inhibitor cocktail (ThermoFisher Scientific; Cat# J61852-XF). Protein lysates were quantified using a Pierce BCA Protein Assay Kit (ThermoFisher Scientific; Cat# 23227), and unless otherwise stated, 50 µg of total whole cell lysate was resolved on 4–20% polyacrylamide Mini-PROTEAN^®^ TGX™ Precast Protein Gels (BioRad). Following an overnight transfer onto PVDF membranes (BioRad; Cat# 1620175), samples were blocked with 5% BSA (RPI Corp.; Cat# A30075) and incubated with primary antibody overnight, washed 3 times, and incubated with secondary antibodies for 1 hour at room temperature. Membranes were washed 4 times for 15 minutes with gentle rocking, before being incubated with SuperSignal™ West Dura Extended Duration Substrate (ThermoFisher Scientific; Cat# 34075). Membranes were imaged on either a ChemiDoc MP Imaging System (BioRad) or a LI-COR Odyssey CLx imager.

### Antibodies

Primary antibodies used in these experiments were validated on dystrophin-deficient tissues or appropriate control tissues for specificity. Primary antisera used for western immunoblotting and immunofluorescent labeling include: dystrophin all isoforms (Abcam; Cat# ab15277), dystrophin Dys1 (Leica; Cat# NCL-DYS1), and vinculin (Cell Signaling Technology; Cat# 7076S). Secondary antibodies used for western immunoblotting included anti-mouse IgG HRP (Cell Signaling Technology; Cat# 7076S) and anti-rabbit IgG HRP (Cell Signaling Technology; Cat# 7074S). Secondary fluorescent anti-mouse and anti-rabbit IgG fluorochrome-conjugated antibodies used include: Alexa Fluor 488 (ThermoFisher Scientific; Cat# A-11001 and A-11008), Alexa Fluor 555 (ThermoFisher Scientific; Cat# A32727 and A32732), and DyLight 633 (ThermoFisher Scientific; Cat# 35563). Fluorescently labeled sections were imaged on a Nikon A1R-HD25 Confocal microscope using NIS Elements software, and later processed using Adobe Photoshop (2025 version).

### Tamoxifen administration

We used a previously-established protocol for the *HSA*-MerCreMer strain of delivering (80 mg/kg body weight) of (Z)-4-hydroxyl-tamoxifen (4-OHT) (MilliporeSigma; Cat# H7904) mixed with sunflower seed oil (MilliporeSigma; Cat# S5007) via 5 daily intraperitoneal injections while alternating left and right sides to avoid injection site reactions (46, 47). Control mice were given equivalent IP injections of sunflower seed oil. Separate mouse cohorts were assessed at 30 day intervals from days 0, 30, 60, 90, and 180 days post injection. Mice were monitored for hepatotoxicity using cytochrome P450 assessments and injection site reactions after administration.

### Mice locomotor and functional analysis

Open field testing and mouse locomotor analysis was measured using a previously established 4 individual chamber sensor system protocol and data was analyzed using Ethovision XT software platform (Noldus; Leesburg, VA)(48, 49). Mice were analyzed for functional analysis but were first acclimated to the room and open-field chambers one day prior to activity and were given a five minute additional adaptation period prior to activity recording. Mouse locomotor activity was recorded for ten minutes with no external stimulation and the experimenter isolated in a separate room.

### Physiological force assessments

Mice were anesthetized with sodium pentobarbital (80 mg/kg) by intraperitoneal (IP) injection. Isolated EDL muscles were suspended in a phosphate buffer equilibrated with 95% O_2_, 5% CO_2_ (35 °C). Muscle contractions were generated using a 150 ms, supramaximal stimulus train (200 µs pulses) with the muscle held at its optimal length (L_o_) for tetanic tension. Force was normalized to physiological cross-sectional area assuming a fiber length to muscle length ratio of 0.44 as previously described(32). After evaluation of peak force, muscles were subjected to an active lengthening protocol consisting of the following sequence: one fix-end contraction, 5 lengthening (eccentric) contractions, and two fixed-end contractions. All contractions began with the muscle at L_o_ and were elicited with 150 ms trains at 300 Hz. The lengthening trials consisted of an initial 100 ms fixed-end contraction that allowed the muscle to rise to peak force followed by a constant velocity stretch at four muscle fiber lengths/s to a final length of 120% L_o_. Force was evaluated at 95 ms of stimulation for both fixed-end and lengthening trials.

### Social neurobehavior and memory experiments

Social approach was determined by placing the test mouse in the middle chamber of a 3-chamber system and determining if the mouse showed preference for a novel mouse or a novel object (round ball). The social approach protocol was adapted from our previous protocol along with other established mouse neurobehavioral assessments in autism spectrum disorder (ASD) mouse models(20, 50). Mice were evaluated over 10 minute periods after a 10 minute acclimation period in which the novel mouse and novel object were kept in blocked off chambers. Placement of the novel mouse and novel object was switched between the two side chambers between intervals, and the apparatus was cleaned between evaluations. Social novelty was determined by placing the test mouse in the center chamber and evaluating the time spent in chambers containing either an unfamiliar mouse (no previous exposure) or a familiar mouse (aged-matched littermate). The mouse was monitored over a 10 min interval after the 10 minute acclimation period, and its total movement, velocity, time spent in close interaction with familiar or unfamiliar mouse were measured. T-maze and Y-maze (Maze Engineers; Skokie, IL) cohort assessments were performed using previously described with video recordings captured after the mouse was placed in the maze center(51, 52). Spatial memory and learning evaluation of the mouse cohorts was performed using the Morris Water Maze (Noldus) from an adapted protocol previously used in *mdx* mice(53, 54). Experimenter was double-blinded to treatment group and genotype of the mice in each cage and de-blinded occurred after data analysis.

### Skeletal muscle injury

A cardiotoxin-induced injury to the TA was performed by intramuscular injection of 40 µl of 10 µM cardiotoxin (Millipore Sigma; Cat# 217503) given to adult, aged-matched male mice. The contralateral TA muscle was used as a sham control and given an equivalent intramuscular injection of phosphate-buffered saline (1x DPBS; Millipore Sigma; Cat# D8662) given as a control. Mice were evaluated at day 0 (uninjured), 7, 14, and 21 days post cardiotoxin injury for histological changes in muscle architecture and myofiber composition.

### RNA-sequencing

Total RNA was extracted from TA skeletal muscles from aged-matched male mice using TRIzol reagent (ThermoFisher Scientific; Cat# 15596026) for extraction following the manufacture’s protocol. RNA with an RNA Integrity Score (RIN) greater than 7.0 was selected for library amplification. Poly A enrichment using a ribominus kit (ThermoFisher Scientific; Cat# K155002) was performed on the samples. A total RNA library was amplified from approximately 200-300 ng of RNA using a NovaSeq 6000 reagent kit v1.5 (Illumina; San Diego, CA; Cat# 20044417) before being processed on an Illumina NovaSeq 6000 sequencer. Total reads were FASTQ files were uploaded to the UAB High Performance Computer cluster for bioinformatics analysis with the following custom pipeline built in the Snakemake workflow system (v5.2.2)(55). Following count normalization, principal component analysis (PCA) was performed, and genes were defined as differentially expressed genes (DEGs) if they passed a statistical cutoff containing an adjusted p-value <0.05 (Benjamini-Hochberg False Discovery Rate (FDR) method) and if they contained an absolute log_2_ fold change >=1. Functional annotation enrichment analysis was performed in the NIH Database for Annotation, Visualization and Integrated Discovery (DAVID, v6.8) by separately submitting upregulated and downregulated DEGs. Ingenuity pathway analysis (IPA) was performed on the data as described(56). A p-value <0.05 cutoff was applied to identify gene ontology (GO) terms and the FASTQ experimental files have been uploaded to NCBI’s Gene Expression Omnibus under accession number GSE284723.

### Quantitative PCR analysis

Total RNA was extracted using TRIzol reagent (ThermoFisher Scientific) and the lysing matrix A beads (MP Biomedicals; Cat# 1169100-CF) as described above. Approximately 1 µg of RNA was then reverse transcribed using SuperScript IV First-Strand Synthesis System (ThermoFisher Scientific; Cat# 18091050). Approximately 0.5 µl of sample was then mixed with using TaqMan™ Fast Advanced Master Mix (ThermoFisher; Cat# 4444556) following the manufacturer’s protocol and samples were analyzed on a QuantStudio 5 Real-Time PCR machine (ThermoFisher). The following Taqman real time PCR assays (ThermoFisher Scientific) were used to amplify specific transcripts: *Dmd* (exon 1, Mm03956078_s1; exons 16-17, Mm00464475_m1; exons 50-51, Mm01216958_m1; exons 64-65, Mm00464520_m1), and *Actb1* (Mm02619580_g1).

### Statistical Analysis

For most dual cohort comparisons, either a two-tailed student’s t-test or chi-squared test for normality was performed for all pairwise comparisons. For longitudinal assessments over time or across multiple cohorts, a one-way analysis of variance (ANOVA) with Least Significant Difference (LSD) was performed for all multiple comparisons. GraphPad Prism version 10 software (Graphpad Software; San Diego, CA) was used for all statistical analyses. An *a priori* hypothesis of *p< 0.05, **p<0.01, ***p<0.001, and ****p< 0.0001 was used for all reported data analyses. All graphs were represented as mean +/- SEM.

## Acknowledgments and Funding

We would like to thank members of the Alexander lab and UAB Center for Exercise Medicine for their critical evaluation of our manuscript. We would also like to thank members of the Kunkel, Gamble, and Esser labs for reviewing our manuscript and assistance with data interpretation. M.S.A. is supported by NIH grants from the Office of the Director under award number (U54OD030167), and an R21 grant from National Institute of Neurological Disorders and Stroke (NINDS) (R21NS140497). Additional funding support was provided by the Hope4Gabe foundation. L.M.K. is supported by NIH National Institute of Arthritis and Musculoskeletal and Skin Diseases (NIAMS) grant R01AR064300, and the Bernard F. and Alva B. Gimbel Foundation. K.L.G. is funded by an NIH National Institute on Drug Abuse (NIDA) grants R01DA046096 and R01NS082413. K.A.E. is funded by NIH NIAMS grants (R01AR066082 and R01AR079220). M.A.L. is supported by an NIH NIAMS grant K08NS120812.

