## SupplementalFigs for "Conditional *Dystrophin* ablation in the skeletal muscle and brain causes profound effects on muscle function, neurobehavior, and extracellular matrix pathways"

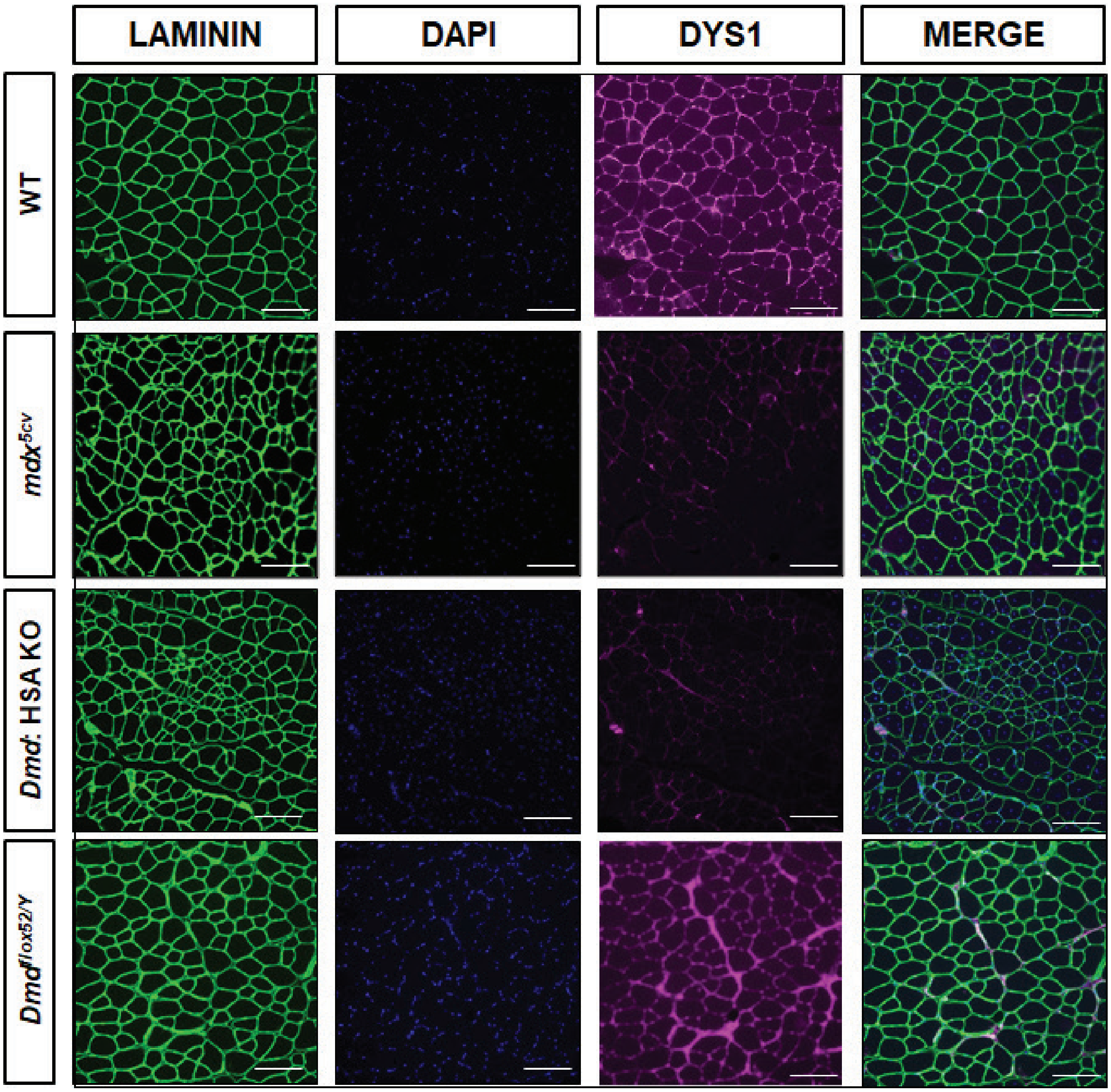


**Supplemental Figure S1: Immunofluorescent confirmation of the loss of skeletal muscle dystrophin.**

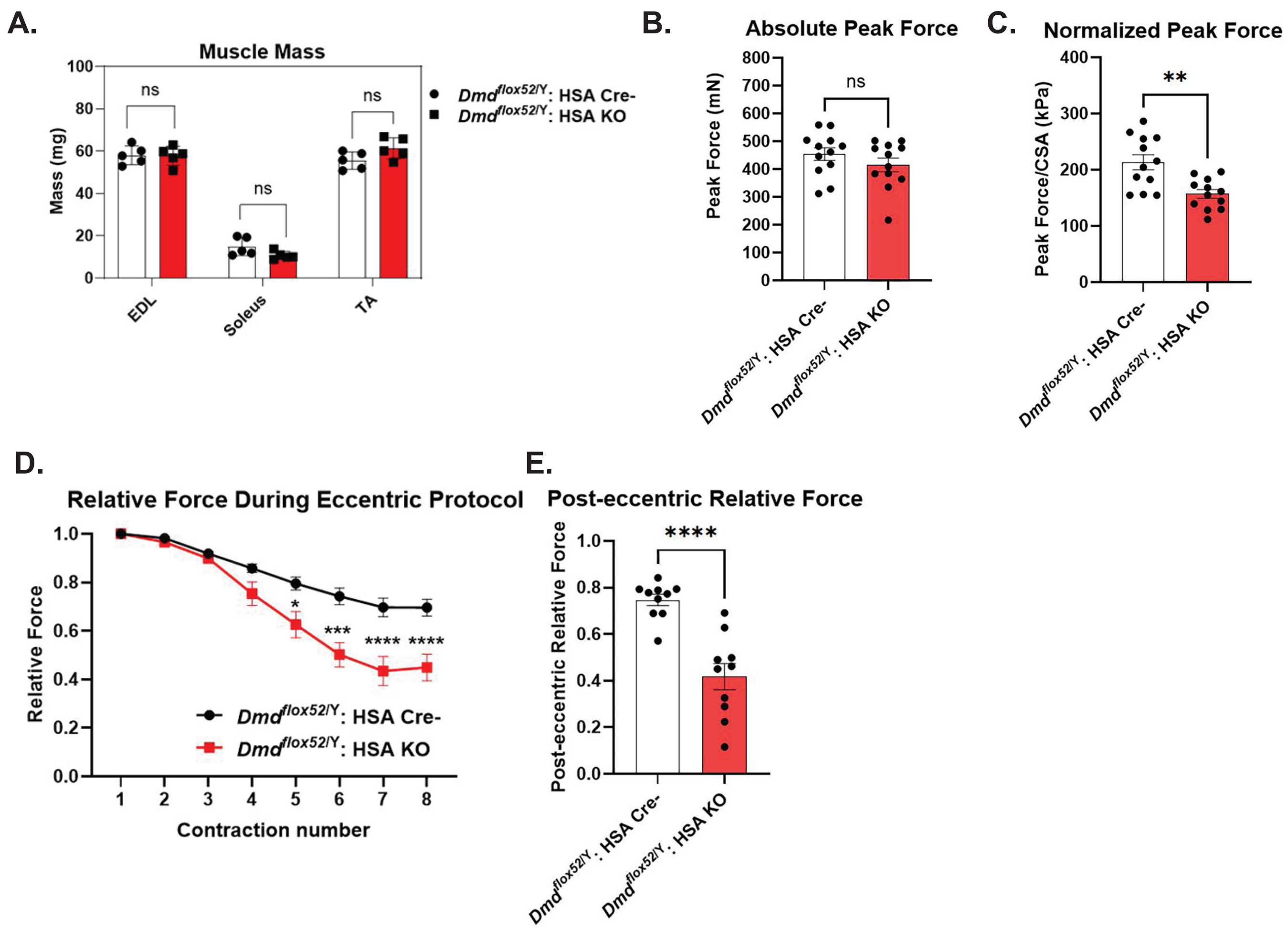


**Supplemental Figure S2: Adult *Dmd* myofiber KO mice have physiological force deficits.**

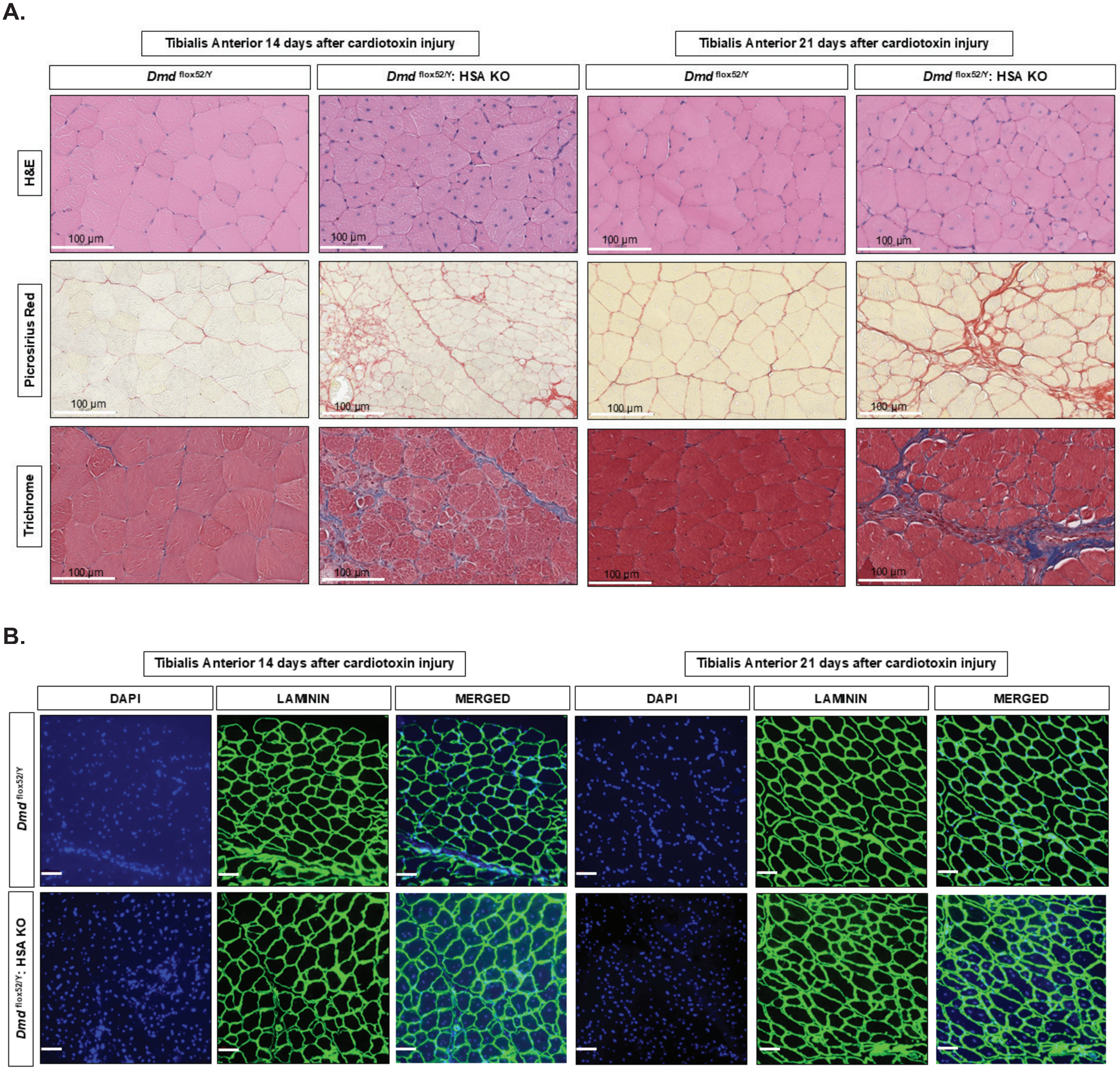


**Supplemental Figure S3: Conditional *Dmd* myofiber knockout mice have impaired skeletal muscle regeneration.**

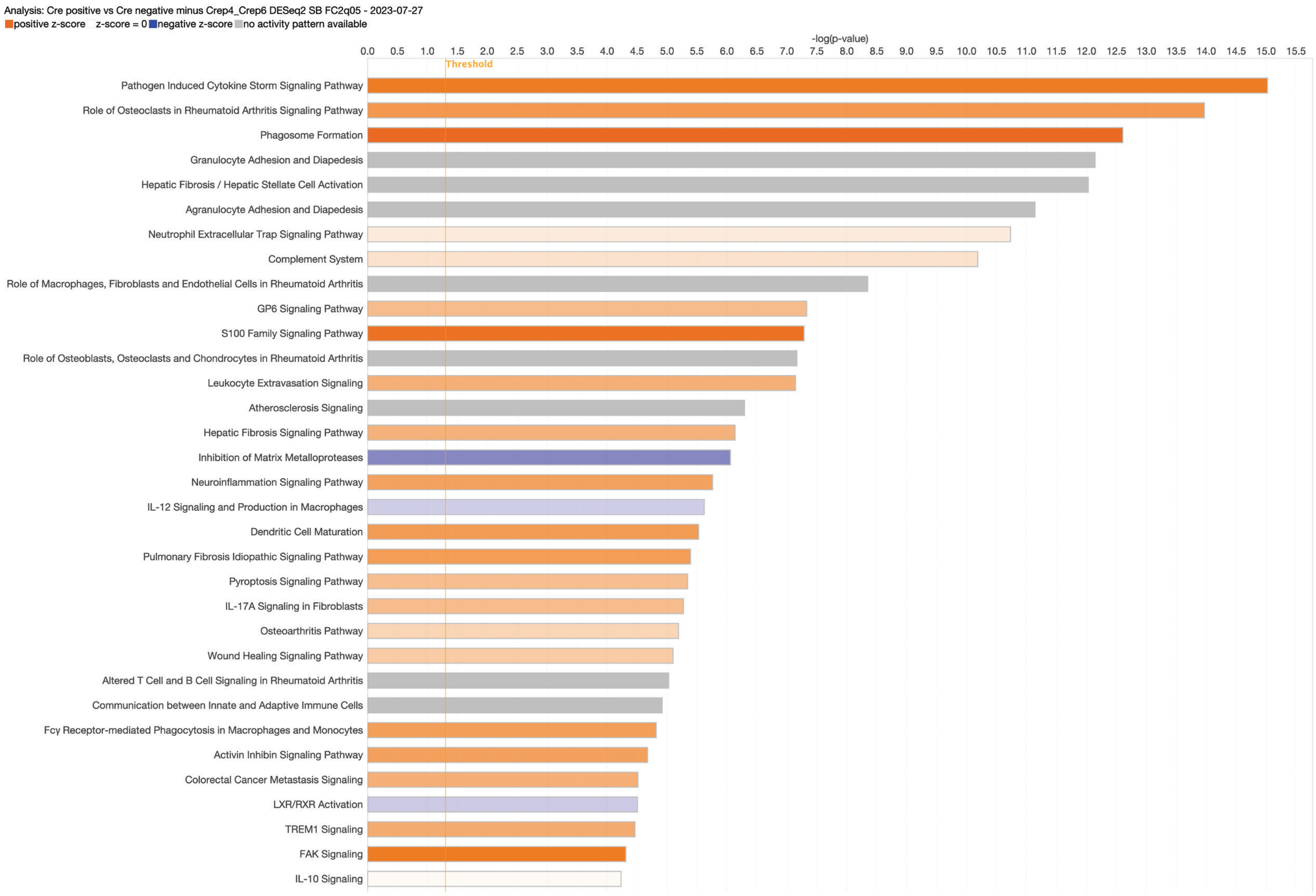


**Supplemental Figure S4: IPA pathway analysis of the conditional *Dmd* myofiber knockout mice.**

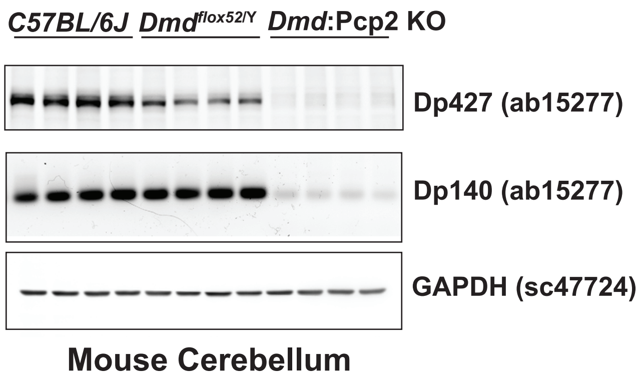


**Supplemental Figure S5: Confirmation of cerebellar dystrophin Dp427 protein knockdown in *Dmd*:Pcp2 isolated cerebellums.**
